# Genome-wide transcript and protein analysis reveals distinct features of aging in the mouse heart

**DOI:** 10.1101/2020.08.28.272260

**Authors:** Isabela Gerdes Gyuricza, Joel M. Chick, Gregory R. Keele, Andrew G. Deighan, Steven C. Munger, Ron Korstanje, Steven P. Gygi, Gary A. Churchill

## Abstract

Investigation of the molecular mechanisms of aging in the human heart is challenging due to confounding factors, such as diet and medications, as well limited access to tissues. The laboratory mouse provides an ideal model to study aging in healthy individuals in a controlled environment. However, previous mouse studies have examined only a narrow range of the genetic variation that shapes individual differences during aging. Here, we analyzed transcriptome and proteome data from hearts of genetically diverse mice at ages 6, 12 and 18 months to characterize molecular changes that occur in the aging heart. Transcripts and proteins reveal distinct biological processes that are altered through the course of natural aging. Transcriptome analysis reveals a scenario of cardiac hypertrophy, fibrosis, and reemergence of fetal gene expression patterns. Proteome analysis reveals changes in energy metabolism and protein homeostasis. We found that for many protein complexes there is a decline in correlation between their component proteins with age, indicating age-related loss of stoichiometry. Some of the most affected complexes are themselves involved in protein homeostasis, which potentially contributes to a viscious cycle of progressive breakdown in protein quality control with age. In addition, we identified genetic loci that modulate age-related changes in a variety of cellular processes, including protein degradation and sorting, suggesting that genetic variation can alter the rate of molecular aging.

## INTRODUCTION

Cardiovascular (CV) diseases are the leading cause of death in elderly people. Improved understanding of mechanisms that underlie the changes that occur in the aging heart could open new opportunities for prevention and treatment (1). As the heart ages, characteristic physiological changes occur, including increased arterial thickening and stiffness, endothelium dysfunction, valvular fibrosis and calcification, and a switch from fatty acid to glucose metabolism (2–4). Compensatory mechanisms may temporarily maintain heart function but can also contribute to progressive deterioration and eventual heart failure (2). For example, thickening of the left ventricle and remodeling of the extracellular matrix may compensate for loss of systolic function (2,3). However, in the long term, the increased wall stress causes the left ventricle to dilate, leading to a decline in systolic function (5). Physiological measures of cardiac function that change with age have high heritability suggesting that genetic factors contribute to variability in cardiac aging in humans (6).

Despite well-known physiological changes in the aging heart, dissecting the cellular and molecular basis of age-related change is challenging due to its complex dynamics and inherent variability in the aging process (7,8). Age-related changes at the cellular levels have been associated with genomic instability, loss of protein homeostasis, epigenetic alterations, mitochondrial dysfunction and inflammation (7). Variability of transcript expression increases with age in mammalian tissues, including the heart (9,10). Age-related dysregulation of transcripts is offset by selective translation, and post-transcriptional mechanisms become crucial for achieving cellular homeostasis (11). The investigation of molecular mechanisms involved in aging is further complicated by the uncoupling of age-related changes between transcripts and their corresponding proteins (12). Waldera-Lupa et al (2014) found that 77% of the proteins that change with age in human fibroblasts showed no corresponding change in their transcripts (13). Decoupling between transcripts and proteins with age has also been observed in the brain of humans and rhesus macaques (14). Thus, investigating age-related changes using only transcriptional profiling may fail to reveal important influences on proteins and higher-order cellular processes.

Mouse models of aging can recapitulate many of the cardiac aging phenotypes seen in humans, such as increased atrial and ventricular dimensions and reduced diastolic function (15), and thus provide relevant models for investigating aging processes in the heart. However, most previous studies have used mice descended from only a few isogenic strains that do not reflect the diversity of cardiac phenotypes found in aging human populations. Multiple studies report differences in mouse cardiac phenotypes, under either physiological or pathological conditions, associated with genetic background across inbred strains (16–20), and in multiparent populations (21,22), confirming the importance of genetic diversity in shaping the rate and course of cardiac aging.

In this study we utilize Diversity Outbred (DO) mice derived from eight inbred founder strains: A/J (AJ), C57BL/6J (B6), 129S1Sv/ImJ (129), NOD/ShiLtJ (NOD), NZO/H1LtJ (NZO), CAST/EiJ (CAST), PWK/PhJ (PWK), and WSB/EiJ (WSB), to investigate cardiac aging in a genetic and phenotypically diverse model (23,24). We analyze transcriptome data from RNA sequencing (RNA-seq) and protein data from mass-spectrometry analysis of heart tissues collected from healthy DO mice at ages 6, 12 and 18 months. At 6 months of age, the mice have reached full maturity. At 18 months, most mice are healthy and are only beginning to show signs of age-related decline. Thus, we are looking at changes in transcripts and proteins that are not influenced by developmental programs and are also not reflecting late-stage disease progression (25). These data can reveal broad patterns of change in biological processes and in specific cellular compartments. To characterize molecular and cellular changes in the aging mouse heart, we first identify the transcripts and proteins that are changing with age and identify functionally related groups of genes using gene-set enrichment analysis (26,27). We then examine the maintenance of protein complex stoichiometry. Finally, we investigate how genetic variation modulates age-related changes in the heart. The molecular profiling data from the mouse heart are freely available to support further investigations of the molecular basis of aging in mammals (https://qtlviewer.jax.org/viewer/agingheart).

## RESULTS

### Transcriptomics reveals age-related changes in muscle cell differentiation, contraction, and inflammation

We used RNA-Seq data from the hearts of 192 DO mice of both sexes aged to 6, 12 or 18 months to identify transcripts that change with age (Figure 1). Transcripts that change with age are referred to throughout as age-related transcripts, and their magnitude and direction of change are referred to as age effects, reported in unit of log_2_ fold change per year. For the transcriptome data, we used DESeq2 (28) to identify 2,287 transcripts (out of 20,932) with significant age-related changes (false discovery rate, FDR < 0.1; Supplementary File 1). These transcripts include genes that are known to play a role in the aging heart, as well as genes that have been implicated in aging but have not been previously reported as changing in the heart, or genes that play a role in heart disease or heart development but have not been reported to change with age. They are enriched for functional annotations across 85 biological processes (FDR < 0.05), the most significant of which are muscle cell migration, regulation of muscle cell differentiation, ion transport pathways, and acute-phase response (Figure 2).

**Figure 1.**
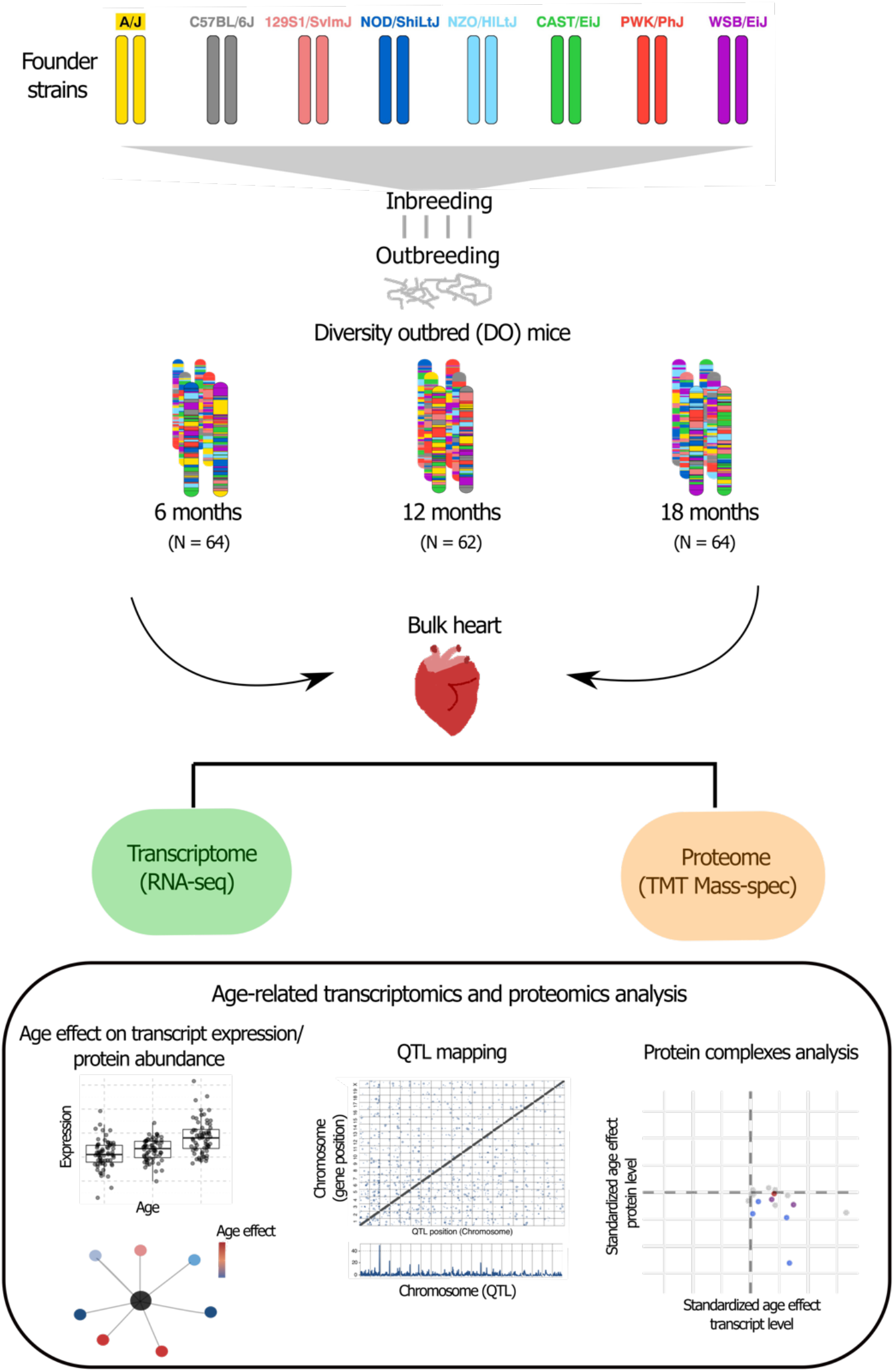
Study Design. We carried out a cross-sectional aging study of DO mice, which are descendant from 8 founder inbred strains and are maintained as an outbred stock with high levels of heterozygosity. Heart tissue from DO mice aged 6, 12 or 18 months was collected for RNA-seq and mass-spectrometry analysis. The transcriptome and proteome data were analyzed to evaluate age-related changes in transcript and protein abundance using gene set enrichment, QTL mapping and protein complex stoichiometry.

**Figure 2.**
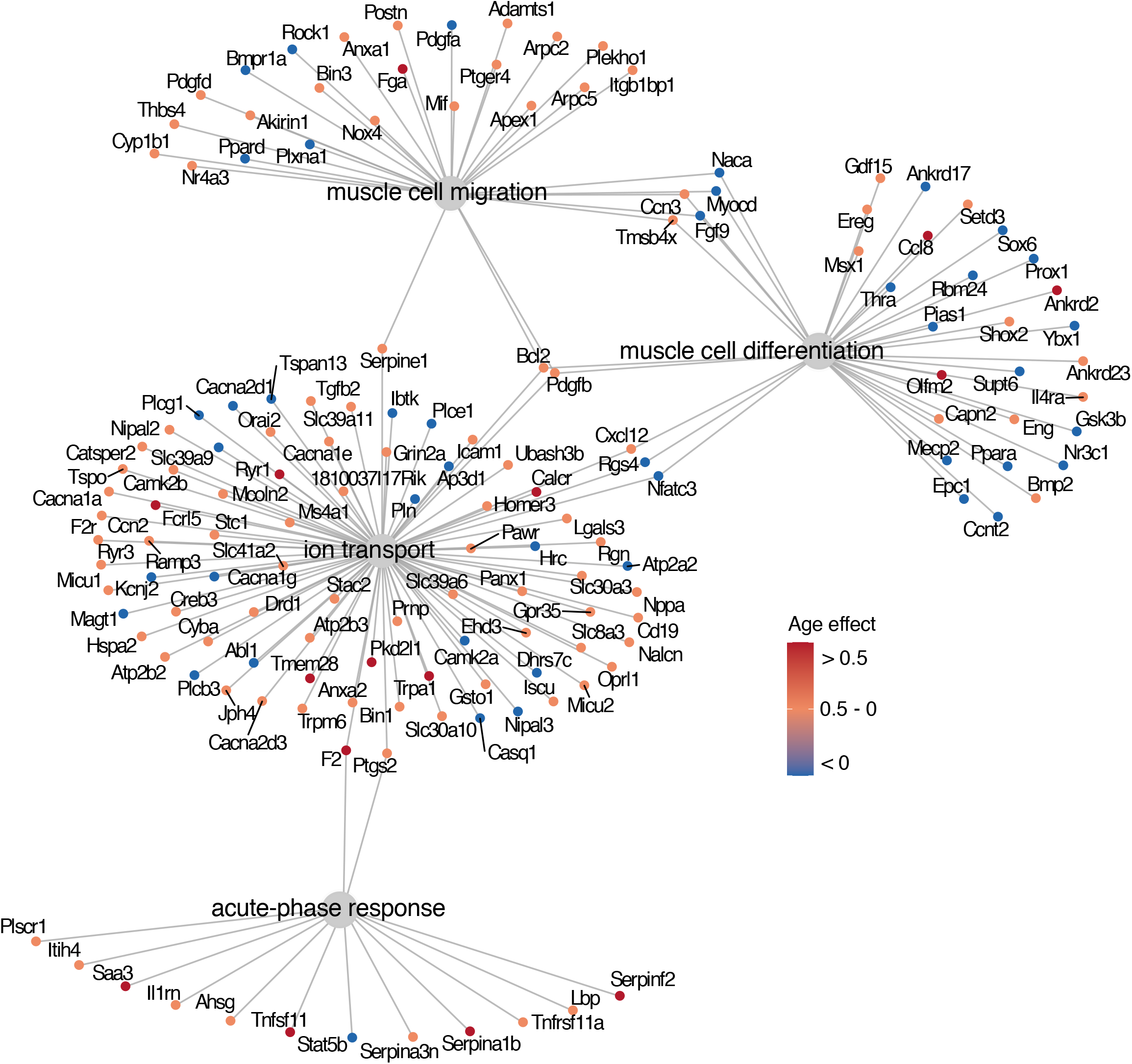
Enriched gene ontology terms for age-related transcripts. The most significant gene ontology categories for age-related transcripts in DO mouse hearts are shown. Gene-set enrichment analysis reveals that age-related transcripts are involved in muscle cell migration, regulation of muscle cell differentiation, ion transport and acute-phase response (FDR < 0.05). Major nodes indicate enriched categories and adjacent nodes identify transcripts within each category. Transcripts involved in muscle cell migration, muscle cell differentiation and ion transport both increase and decrease with age. Transcripts from the acute-phase response increase except for *Stat5b*. The color scale represents the age effect for each transcript in units of log_2_ fold change (LFC) per year.

Some of the age-related transcripts possess functions relevant to cardiac pathological conditions and heart development (Table 1). Both *Adamts1*, present in the muscle cell migration pathway, and *Serpine1*, present in both muscle cell migration and ion transport pathways, increase with age (*Adamts1*: age effect = 0.15; *Serpine1*: age effect = 0.19; Figure 3A). These genes have been shown to induce collagen 1 deposition and fibrosis in the heart (29–31). The increased expression of transcripts involved in fibrosis is indicative of age-related changes in cellular composition including an increase in the proportion of fibroblasts and myofibroblasts in the heart. Comparing the age-related transcripts reported here with published single-cell RNA-seq data (18), we observe an increase in expression of fibroblast markers including *Timp1* (age effect = 0.3) and *Mt2* (age effect = 0.4), and an increase in myofibroblast markers including *Postn* (age effect = 0.3) and *Cthrc1* (age effect = 0.46). We also observed increased expression of *Nov* (age effect = 0.18; Figure 3A), which plays a role in heart development by blocking terminal differentiation and increasing the proliferation rate of myoblasts (34, 35).

**Table 1.** Highlighted transcripts that change with age.

| Transcript symbol | Direction of change | Function highlights | Reference index |
| --- | --- | --- | --- |
| *Adamts1* | Increases | Cardiac fibrosis | (29) |
| *Ahsg* | Increases | Free fatty-acid signaling | (43) |
| *Nov* | Increases | Heart development | (32,33) |
| *Olfm2* | Increases | Vascular smooth muscle cell regulation | (126) |
| *Serpine1* | Increases | Cardiac fibrosis | (30,31) |
| *Tnfsf11* | Increases | Aortic valve calcification | (45) |
| *Cacna1g* | Decreases | Cardiac muscle contraction | (127,128) |
| *Myocd* | Decreases | Cardiomyocytes survival | (34) |
| *Naca* | Decreases | Sarcomere organization | (129–131) |
| *Pln* | Decreases | Cardiac muscle contraction | (37) |
| *Ppara* | Decreases | Fatty acid metabolism | (132–134) |
| *Stat5b* | Decreases | Insulin signaling | (40,41) |

**Figure 3.**
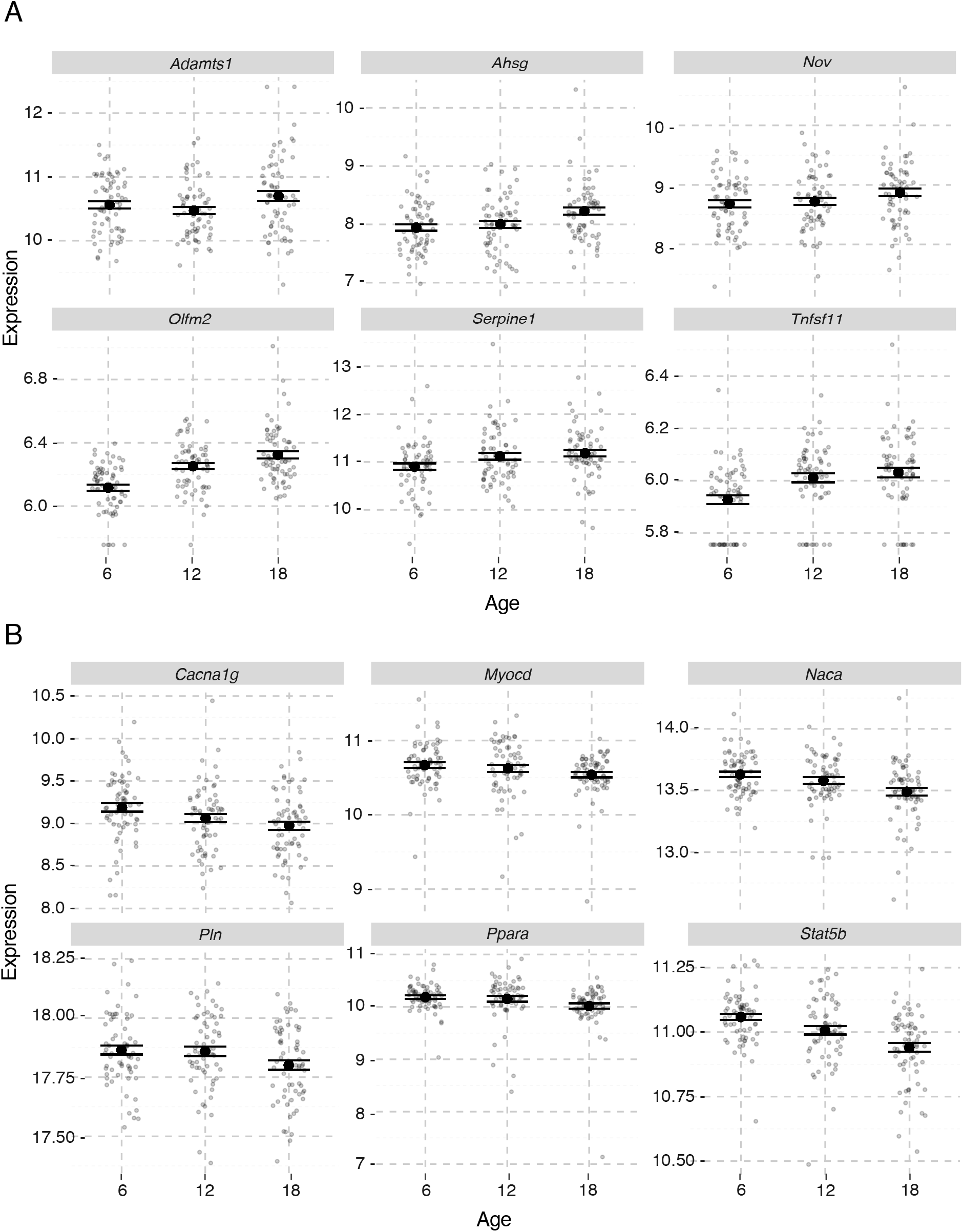
Individual-level expression of age-related transcripts. Transcripts highlighted in Table 1 are shown across age groups (6, 12 and 18 months). Transcripts increase (A) or decrease (B) with age. Solid points represent the age group mean expression +/- 1 standard error (SE), and points with transparency show individual mouse expression levels. Expression levels (Y axis) are counts normalized by the DESeq2 variance stabilizing transformation.

The gene *Myocd*, whose expression decreases with age (age effect = -0.12; Figure 3B), is involved in both muscle cell migration and muscle cell differentiation, and it is an important regulator of cardiac function by maintaining cardiomyocyte cell structure and function (34). *Ppara* also decreases with age (age effect = -0.11 – Figure 3B), and it plays a role in the regulation of cardiac fatty acid metabolism and is implicated in several pathologic heart conditions associated with aging (35,36). Many age-related transcripts that are associated with calcium ion transport, including *Pln, Cacna1g* and *Dhrs7c*, decrease with age (Table 1 and Figure 3B). These genes have functions associated with cardiac muscle contraction, play a role in cardiomyopathy, and are downregulated in heart failure models but have not been previously described in the aging heart (37–39).

Acute phase response pathway genes, which are involved in inflammation, increase with age except for *Stat5b*, which decreases (age effect = -0.07; Figure 3B). Although its function in heart is not established, STAT5B interacts with the insulin receptor, coordinating changes in gene expression through insulin signaling (40,41). In addition, STAT5B was proposed to inhibit acute-phase response by modulating the activation of STAT3 (42). Transcripts in the acute phase response pathway that increase with age include *Ahsg* (age effect = 0.3; Figure 3A), which controls the binding of free fatty-acid to inflammatory receptors and protects against vascular calcification (43,44), and *Tnfsf11* (age effect = 0.52 – Figure 3A), which is involved in aortic valve calcification in response to inflammation, a feature prevalent in the elderly (45). We observe an age-related increase in expression of immune cell-specific markers, such as B cells (*Cd79a*, age effect = 0.46), macrophages (*Cd68*, age effect = 0.19) and monocytes (*Plac8*, age effect = 0.6), indicating immune cell infiltration in the aging heart. The increase in expression of immune response genes with age in multiple tissues of the mouse, including heart, has been reported previously (46).

### Proteomics reveals age-related changes in mitochondrial metabolism and intracellular protein transport

We obtained proteomics data from 190 of the 192 DO mice that had transcriptome data (Figure 1). In order to test for age-related changes at the protein level, we fit a regression model of protein abundances with a linear term for age and covariates as described in Methods. We identified 1,161 age-related proteins (out of 4,062; FDR < 0.05; Supplementary File 2). A higher proportion of proteins exhibited change with age when compared to transcripts, and thus we applied a more stringent FDR threshold to focus on the proteins with greatest change. These proteins are enriched (FDR < 0.05) for gene ontology categories that include positive regulation of cellular proliferation, intracellular protein transport and several mitochondrial categories (Figure 4; Table 2). Changes in the mitochondrial respiratory chain complexes were also reported in recent studies of single-cell RNA-seq and proteome data of multiple tissues in aging mice (46,47). Interestingly, there was no overlap of enrichment terms for proteins and those found in our transcriptome analysis. The proteome data also highlight different cell-specific markers (18). In addition to fibroblast markers, we see an increase with age in markers of smooth muscle (VTN, age effect = 0.58), epicardium (CLU, age effect = 0.38) and endothelium cells (FABP4, age effect = 0.39 and PECAM1, age effect = 0.3) that were not seen in the transcriptome data. Thus, the protein data reveal unique features aging that are not seen at the transcript level.

**Table 2.** Highlighted proteins that change with age.

| Protein symbol | Direction of change | Function highlights | Reference index |
| --- | --- | --- | --- |
| AKT1 | Increases | Physiological cardiac growth | (48,50) |
| AKT2 | Increases | Cardiac glucose metabolism | (48,51) |
| COX4I1 | Increases | Mitochondrial respiratory chain complex IV cytochrome c subunit | (135) |
| NDUFB6 | Increases | Mitochondrial respiratory chain complex I subunit | (136,137) |
| RABGEF1 | Increases | Mitophagy induction | (56) |
| SDHD | Increases | Mitochondrial respiratory chain complex III cytochrome b subunit | (138) |
| ACTN3 | Decreases | Muscle anaerobic metabolism | (59) |
| ATP1F1 | Decreases | Inhibition of mitochondrial ATPase activity | (139,140) |
| CRIP2 | Decreases | Smooth muscle tissue differentiation and cardiomyocyte survival | (141–143) |
| NDUFA1 | Decreases | Mitochondrial Respiratory Chain Complex I subunit | (144) |
| RAB1B | Decreases | Cardiac hypertrophy | (145) |
| TIMM29 | Decreases | Translocase of inner mitochondrial membrane (Complex 22) | (146) |

**Figure 4.**
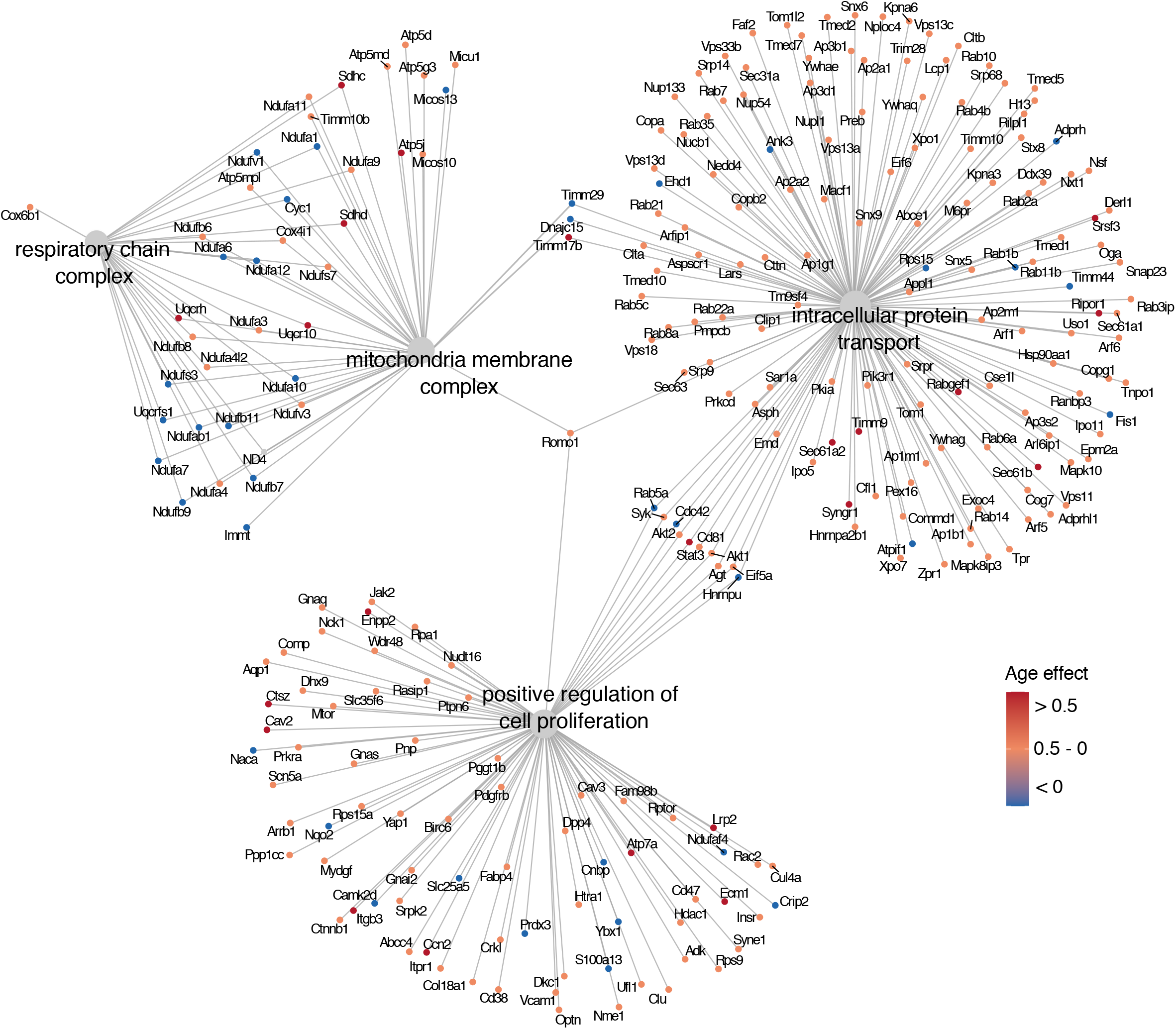
Enriched gene ontology terms for age-related proteins. The most significant gene ontology categories for age-related proteins in DO mouse hearts. Gene-set enrichment analysis reveals that proteins changing with age are involved in positive regulation of cell proliferation, intracellular protein transport, mitochondria membrane complex and respiratory chain complex (FDR < 0.05). Major nodes indicate enriched gene ontology terms and adjacent nodes identify age-related proteins within each category. Most age-related proteins increase in abundance with age, including proteins involved in intracellular protein transport and positive regulation of cell proliferation. Color scale represents the age effect in units of LFC per year.

The serine/threonine kinases AKT1 and AKT2 are highly abundant in cardiomyocytes and regulate cellular proliferation and intracellular protein transport pathways. Both are significantly increasing in abundance with age (AKT1: age effect = 0.36; AKT2: age effect = 0.27 – Figure 5A). AKT1 and AKT2 respond differently to growth factors and extracellular ligands (48), but they both participate in the regulation of cardiac hypertrophy in aging through interaction with Sirtuins (49). Knockout mouse models have shown that the lack of AKT1 constrains the ability of cardiomyocytes to respond to physiological hypertrophy. Alternatively, AKT2 mutant mice showed reduced glucose oxidation in heart cells, but normal response to exercise-induce hypertrophy, demonstrating that these two proteins regulate heart remodeling in response to stress in distinct ways (48,50,51). Along with AKT2, other proteins associated with fetal metabolism increase with age, including ACACB (age effect = 0.14), which inhibits fatty acid oxidation, and GYS1 (age effect = 0.23), which plays a role in glucose metabolism (50,52,53).

**Figure 5.**
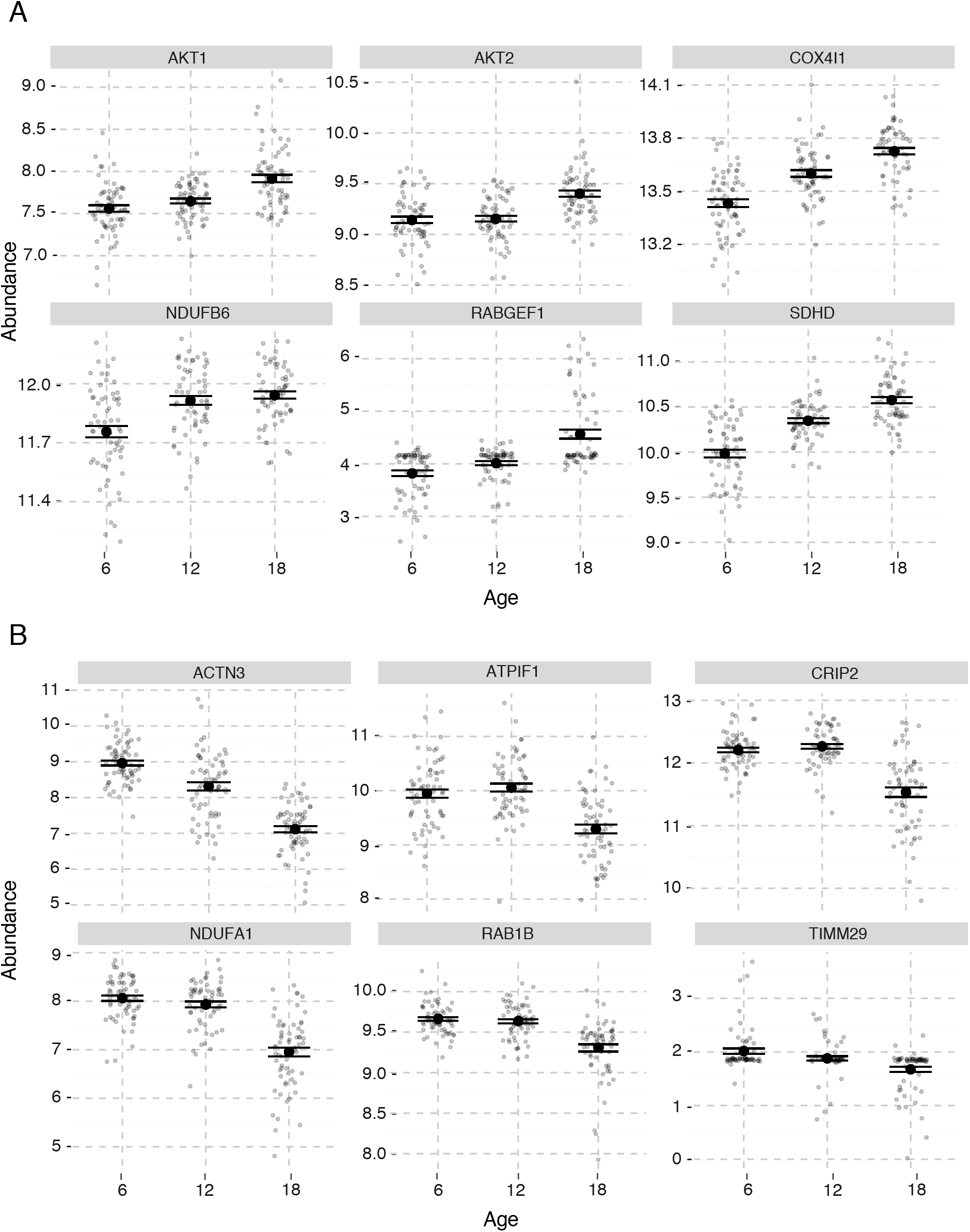
Individual-level expression of age-related proteins. Abundance of proteins highlighted in Table 2 across age groups (6, 12 and 18 months) shows proteins that increase (A) or decrease (B) with age. Solid points represent the age group mean expression +/- 1 standard error (SE), and points with transparency show individual mouse abundance. Abundances (Y axis) are normalized log_2_ values from mass-spec quantification (Methods).

RAB family proteins are Ras-like GTPases that regulate protein trafficking by vesicle formation and fusion throughout the cell (54,55). Among the 22 RAB proteins that change with age, 19 are increasing (Supplementary File 2). Some RAB proteins are activated during mitophagy, which is mediated by RABGEF1 in mammalian cultured cells (56). Notably, RABGEF1 is the RAB family member with the greatest age-related increase (age effect = 0.74; Figure 5A), suggesting that mitophagy may play a substantial role in the aging heart. Increased myocardial RAB abundance is associated with myocardial hypertrophy. Mice overexpressing RAB1A showed contractile depression with impaired calcium reuptake and developed hypertrophy that progressed to heart failure (54). RAB1A and RAB1B are mostly identical in structure and function (57) and RAB1B is one of the few RAB proteins that decreased with age in our dataset (age effect = -0.356; Figure 5B).

We observed distinct patterns of age-related changes in proteins of the mitochondrial respiratory complexes (Figure 5 - Figure supplement 1). Among 16 proteins in the cytochrome C oxidase (COX) complex, i.e., mitochondrial respiratory chain complex IV, 7 show significant age-related changes and, of those, 6 increase in abundance with age. The exception is ACTN3 (alpha-actinin-3), which was decreasing with age (age effect = -1.8; Figure 5B). A role for ACTN3 has not been described in the heart, but in skeletal muscle, it is known to be a negative regulator of oxidative metabolism. Depletion of ACTN3 results in overexpression of COX proteins and to higher mitochondrial oxidative metabolism (58,59). Our findings suggest that there is an increase in oxidative mitochondrial metabolism in the aging heart (Figure 5A; Figure 5 – Figure supplement 1). Most proteins from the SDH and ATP families (succinate dehydrogenase complex – Complex II and ATP synthase complex – Complex V, respectively) increase with age (Figure 5A; Figure 5 – Figure supplement 1). In contrast, subunits from the mitochondrial complex I, i.e. NDUF family (NADH: ubiquinone oxidoreductase supernumerary subunits – Complex I), change in both directions with age (Figure 5A; Figure 5B; Figure 5 – Figure supplement 1).

### Dysregulation of protein complex stoichiometry with age

Loss of stoichiometry in protein complexes has been shown to occur with age in a number of organisms (60–63). We examined the balance of gene-pairs, for both transcript expression and protein abundance, in the mouse heart for 16 complexes as defined in the CORUM database (64). We selected complexes with transcripts and proteins present in our data, including 26S proteasome, nuclear pore complex, cytoplasmic ribosomal small subunit, cytoplasmic ribosomal large subunit, mitochondrial ribosomal large subunit, coat protein I (COPI) vesicle transport, coat protein II (COPII) vesicle transport, mitochondrial ribosomal small subunit, mitochondrial respiratory chain complexes (I-V), mitochondrial pyruvate dehydrogenase complex, mitochondrial inner membrane presequence translocase complex, and mitochondrial outer membrane translocase complex. We computed the Pearson correlations between all pairs of proteins and among all pairs of transcripts within each complex and estimated the age-related changes in correlation (Methods). We evaluated the significance of these changes using a permutation procedure (65).

We identified 123 protein-pairs (out of 2074) with significant changes in correlation with age (FDR < 0.1) (Supplementary File 3). Of these, 4 are in the cytoplasmic ribosomal small subunit complex, 11 are in the COPII complex, 27 are from the mitochondrial respiratory chain complexes (5 from mitochondrial complex I, 2 from mitochondrial complex II, 5 from mitochondrial complex III, 4 from mitochondrial complex IV and 11 from the mitochondrial complex V), and 81 are in the 26S proteasome complex. The majority (115 out of 123 pairs) of the protein-pair correlations decrease with age, consistent with the expected loss of stoichiometric balance in these complexes.

As mentioned above, changes associated with the 26S proteasome complex were outstanding, specially at the protein level. For individual protein pairs, the age effects (change in correlation per year) for all the gene-pairs, including the non-significant ones, range from 0.37 to -0.73, while for their transcripts, the range is 0.56 to -0.5 (not significant; Figure 6A-D). The 26S proteasome complex is composed of two subcomplexes, a core particle (20S proteasome) and a regulatory particle (19S proteasome). The core particle of the proteasome is made up of α and β subunits (proteins from the PSMA or PSMB families), while the regulatory particle includes proteins from the PSMC and PSMD families (66). Even though both subcomplexes show significant changes in correlation with age, PSMD14, from the regulatory particle, was found in the greatest number of protein-pairs that change in correlation with age (14), followed by proteins PSMD13 (10) and PSMD11 (9). This suggests that the age-related changes mostly affect the regulatory particle of the proteasome (Figure 6D2).

**Figure 6.**
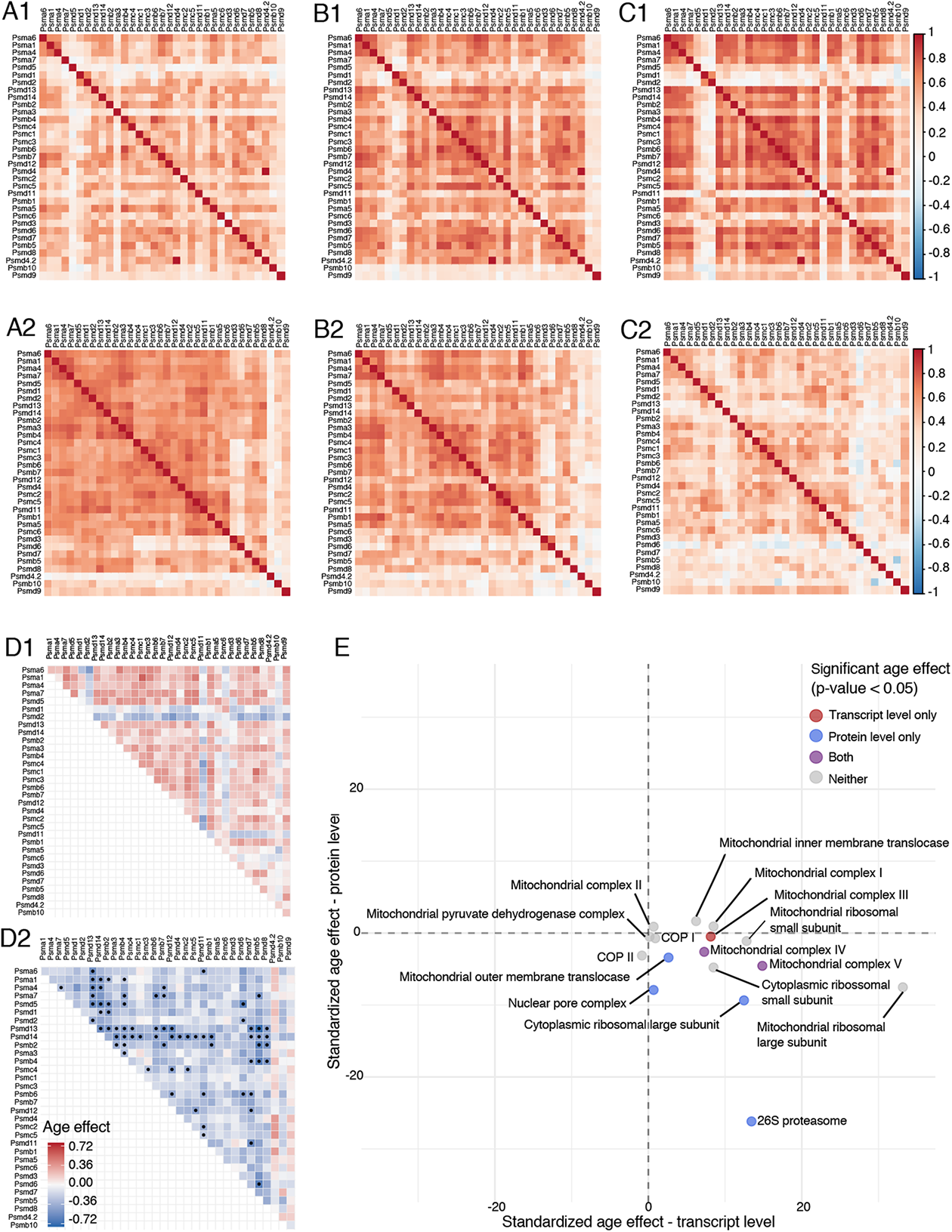
Correlation between protein complex members changes with age. Heatmaps show the correlation coefficients between gene-pairs in the 26S proteasome complex at 6 months (A), 12 months (B) and 18 months (C) at the transcript level (1) and at the protein level (2). D) Heatmaps show the change in correlations between members of the 26S proteasome complex. Correlations among transcripts are generally increasing with age but the change is not statistically significant (D1). The majority of correlations among proteins (D2) are decreasing with age. Dots on the heatmap indicate significant changes in correlations with age (FDR < 0.1). E) Standardized age effects on correlation for 16 protein complexes are shown for transcripts (x-axis) and proteins (y-axis). Color indicates the significance of the age effect on transcripts, proteins, both or neither. Even though many of the complexes do not show a significant change with age, the direction of changes tends to be increasing for transcripts and decreasing for proteins. Age effects are estimated by linear regression and reported as change in the correlation coefficient per year.

Among transcripts, only 20 pairs (out of 2042) showed significant correlation change with age (FDR < 0.1). All of these are in mitochondrial complexes (3 from mitochondrial complex III, 15 from mitochondrial complex V, and 2 pairs from the mitochondrial pyruvate dehydrogenase complex). Interestingly, all of these transcript-pairs increase in correlation with age (Supplementary File 3). Only two of these pairs also showed a significant change at the protein level (*Atp5*b – *Atp5a1* Mitochondrial complex V: protein age effect = -0.40; transcript age effect = 0.21, and *Uqcrc2* – *Uqcrh* Mitochondrial complex III: protein age effect = 0.38; transcript age effect = 0.30).

As an alternative approach to assess coordinated changes across an entire protein complex, we fit a joint model to each set of within-complex correlations with a random intercept term for each gene-pair and a common slope to capture the average correlation change for the complex (Methods). We applied the same permutation test to evaluate statistical significance, but because there is one test per complex, we applied a nominal significance threshold (*p* < 0.05) per complex. Mitochondrial complex III showed an overall change in correlation at the transcript level (age effect = 0.3). We observed changes in only protein correlations for the mitochondrial outer membrane translocase (age effect = -0.21), nuclear pore complex (age effect = -0.16), cytoplasmic ribosomal large subunit (age effect = -0.1) and 26S proteasome (protein age effect = -0.2) (Figure 6E). Mitochondrial complexes IV and V showed change in correlation for both transcripts and proteins (transcript age effect ∼ 0.2; protein age effect ∼ -0.15). We plotted the age effects at the transcript and protein levels for all 16 complexes (Figure 6E), and note that most complexes, regardless of statistical significance, show an increase in transcript correlations and a decrease in protein correlations. Our findings are consistent with previous reports of protein complex stoichiometry changes reported in killifish brain (62), where protein complexes showing the greatest change with age include the mitochondrial complexes IV and V, the cytoplasmic ribosome, and the 26S proteasome complex.

### Genetic variants alter the age trajectory of functionally related groups of proteins

We carried out genetic mapping analysis of transcripts and proteins to identify quantitative trait loci (eQTL and pQTL, respectively) that regulate their expression/abundance levels. The additive effects of genetic variation on transcripts and proteins have been widely documented (67–71) and will not be discussed here. Our interest is to investigate how genetic variants influence the rate or direction of change with age of transcripts and proteins. To identify these age-interactive QTL (age-QTL), we evaluate an age-by-genotype interaction term in a linear mixed model of genetic effects (Methods) for each transcript and protein. We computed genome-wide adjusted significance using permutation analysis (Methods) and declare age-QTL when the age-interactive LOD score (LOD_int_) > 7.75. We have provided access to the data and webtool (QTLviewer) that can be used to explore both the additive and age-interactive genetic effects on transcripts and proteins in the aging mouse heart (https://qtlviewer.jax.org/agingheart). A users’ guide for the QTLviewer can also be found at this website (https://qtlviewer.jax.org/userguide).

We found 824 transcript age-QTLs (age-eQTL; Figure 7A; Supplementary File 4). Most age-eQTL mapped to locations that are distant from their coding genes, which suggests that genetic modification of age-related changes is largely not due to direct regulation of gene expression but rather occur in response to other age-related changes. There are, however, five local age-eQTL. A local age-eQTL for the gene *Cluh* (LOD_int_ = 7.9) is located on chromosome 12. This gene binds RNAs of nuclear-encoded mitochondrial proteins, regulating mitochondrial metabolism by translation control and mRNA decay (72,73). *Cluh* orchestrates mitochondrial metabolic switching from glycolysis to oxidative phosphorylation that happens after birth (73). Genetic variation in *Cluh* might influence the return to the fetal metabolism in the aging heart.

**Figure 7.**
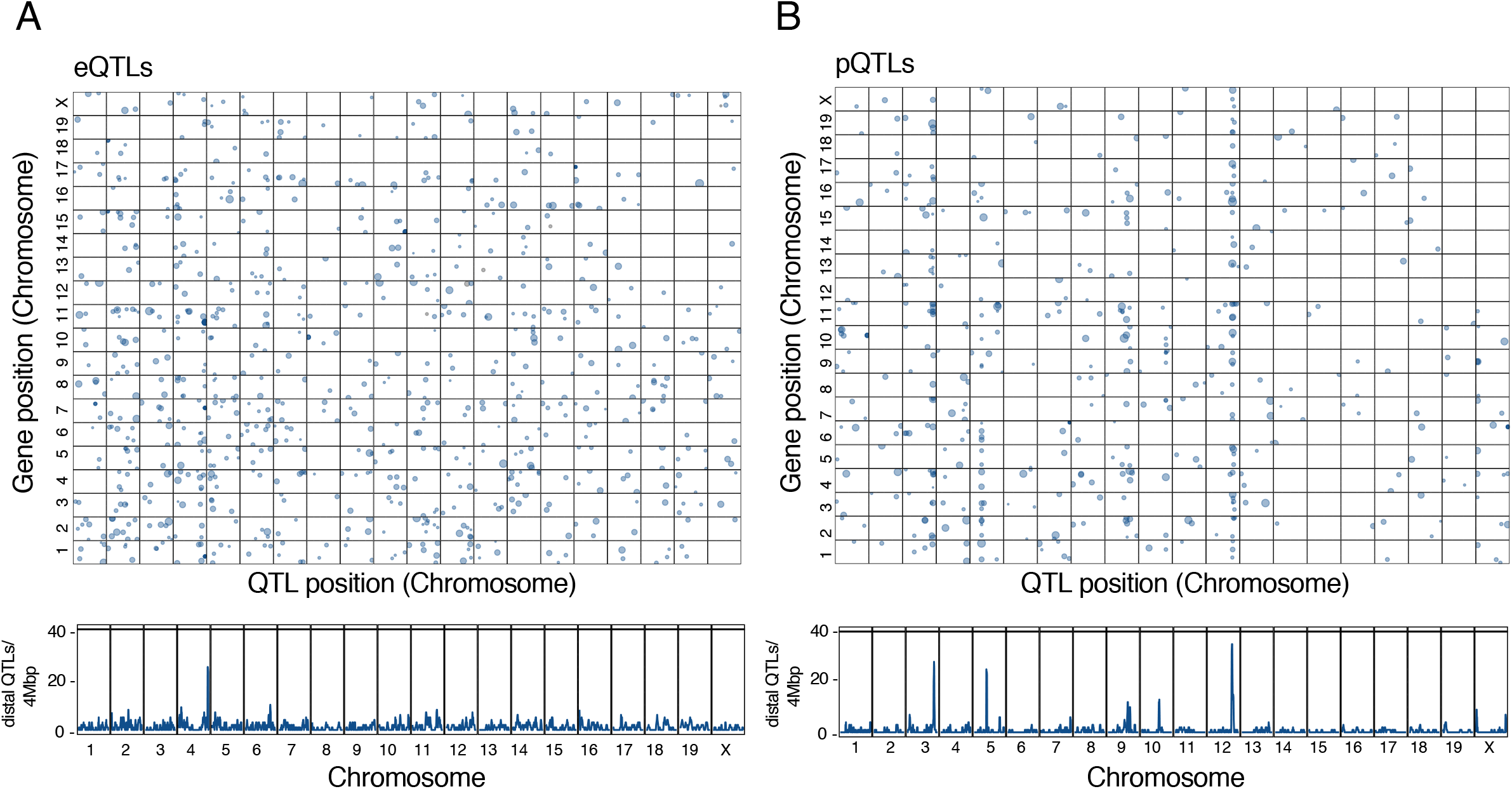
Genomic hotspots for age-interactive eQTL and pQTL. Age-QTL identify the genetic loci involved in age-by-genotype interactions that alter the aging trajectories of transcript (A) or protein (B). In the upper panels in A) and B) points indicate the location of the QTL peak (x-axis) and the location of the coding gene (y-axis) for significant age-QTL (LOD_int_ > 7.75). The lower panels show the number of age-QTL detected in 4Mb windows along the genome. Most age-QTL are distant from the coding gene. Age-QTL can occur in clusters or hotspots where a single genetic locus interacts with many transcripts or proteins. Hotspots can be seen as vertical bands in the upper panels and as peaks in the density plot of age-QTL shown in the lower panels.

We found 463 protein age-QTLs (age-pQTL; Figure 7B; Supplementary File 4), all of which are distant from the coding gene. The protein PARK7, which mapped to a distal age-pQTL on chromosome 9 (LOD_int_ = 10.5) at ∼95Mb, is a redox-sensitive chaperon that protects the murine heart from oxidative damage (74). SIRT2, another protein with relevant function to the aging heart, has a distal age-pQTL on chromosome 13 at ∼111Mb (LOD_int_ = 9.6). SIRT2 regulates cardiac homeostasis and remodeling through activation of AMP-activated protein kinase (AMPK) and repression of the nuclear factor of activated T-cells (NFAT), playing a protective role in the heart (75,76).

Many of the age-QTL co-locate to the genome in hotspots (Figure 7). We identified an age-eQTL hotspot on chromosome 4 and three age-pQTL hotspots on chromosomes 3, 5 and 12, respectively (Methods). Genome-wide significant age-QTL meet stringent statistical criteria that can result in missing weaker but biologically relevant age-QTL. Therefore, at each hotspot, we also considered transcripts or proteins with suggestive QTL (LOD_int_ > 6). We then computed the correlation of all candidate transcripts or proteins within each hotspot and retained only those with absolute mean correlation greater than 0.3. This filter removed genes with age-QTL that are not tightly correlated with other genes at the hotspot and thus less likely to share common genetic regulators. For the age-eQTL hotspot on chromosomes 4, none of the transcripts met this criterion suggesting that the hotspot genes are regulated by multiple independent genetic variants. However, all three age-pQTL hotspots included highly correlated proteins, with 167 at the chromosome 3 locus, 130 at the chromosome 5 locus, and 177 proteins at the chromosome 12 locus (Figure 7 – figure supplement 1; Supplementary File 4).

To determine if the proteins that map to the age-pQTL hotspots share common biological functions, we performed enrichment analysis. The proteins in the chromosome 3 hotspot are enriched (FDR < 0.05) for genes in the proteasome complex, including one protein from the core particle (PSMA family) and several proteins from the regulatory particle (PSMC and PSMD families), including PSDM11 and PSMD14, which are the proteins most affected by age in our correlation analysis (Figure 6D2). Proteins that mapped to the hotspot on chromosome 3 are also involved in the myosin filament, including MYH6 and MYH3 that are responsible for muscle cell structure; and the nucleosome, which is composed of the histone families H1, H2 and H3 (Figure 8A). The proteins that mapped to the chromosome 5 hotspot are associated with 24 enriched GO categories (FDR < 0.05). The two most significant are the endoplasmic reticulum lumen, composed of proteins of the endoplasmic reticulum stress response, such as PDIA4, HSPA5 and HSP9B1, and the contractile fiber containing proteins associated with the muscle contraction apparatus, such as Titin and Desmin (Figure 8B). Finally, for the chromosome 12 hotspot, we found more than 40 GO categories (FDR < 0.05), the most significant of which is muscle contraction that includes proteins related to muscle cell organization, such as myosins and actin, as well as proteins involved in calcium transportation (CACNA2D1) and mitochondria energy generation (NDUFS6) (Figure 8C). Another significant category at the chromosome 12 hotspot is non-coding RNA (ncRNA) transcription, which is associated with chromatin organization, and includes histone H2 family members and a histone chaperone (NPM1) (Figure 8C).

**Figure 8.**
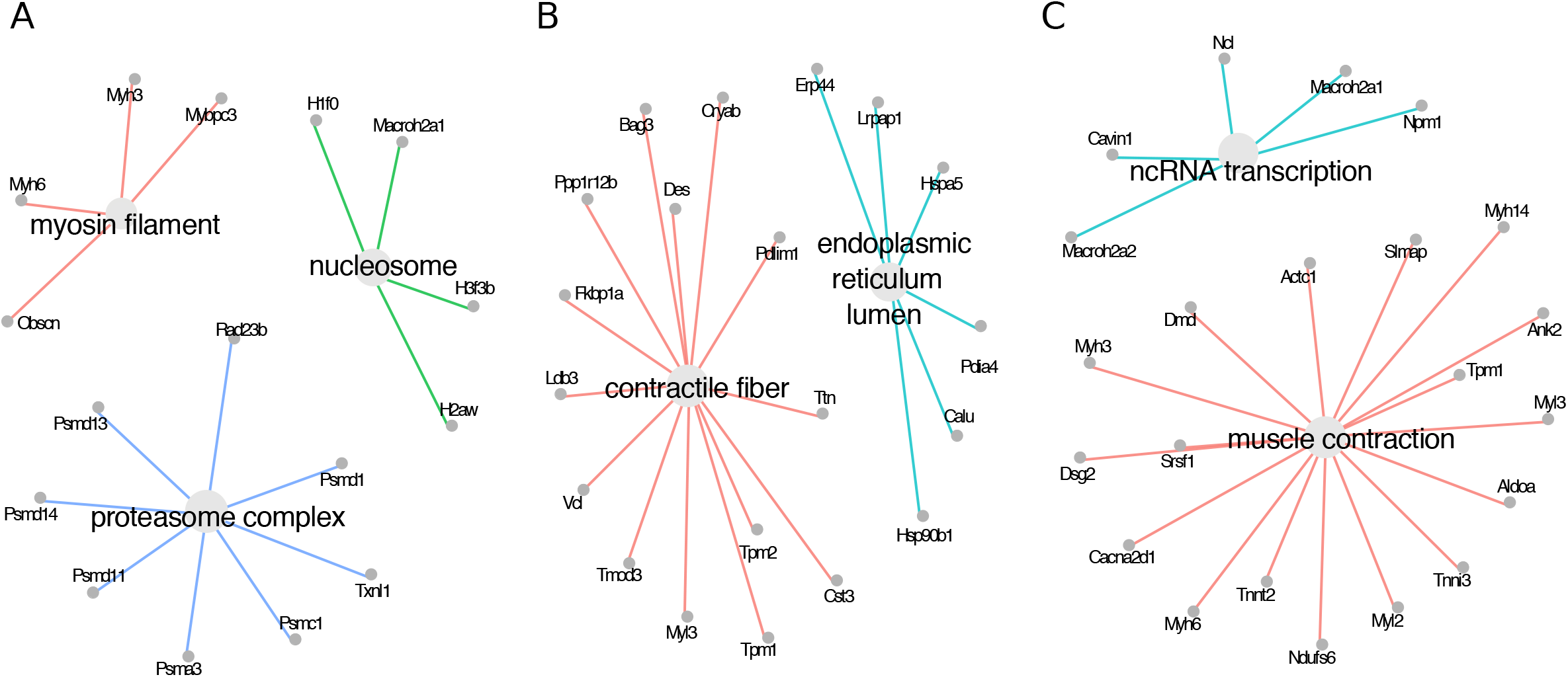
Enriched pathways for proteins in the age-pQTL hotspots. Top enriched GO categories for proteins mapping to the age-pQTLs hotspots are shown. The three hotspot regions regulate proteins involved in distinct biological processes. Proteins that map to the chromosome 3 hotspot (A) are primarily associated with myosin filament, nucleosome and proteasome complex, while proteins that map to the chromosome 5 hotspot (B) participate in contractile fiber and endoplasmic reticulum lumen. Proteins that map to the chromosome 12 hotspot (C) are associated with muscle contraction and ncRNA-transcription.

To investigate how changes in protein abundance with age can vary due to genetic factors, we estimated the age-effects attributable to each DO founder allele by fitting a regression of the first principal component (PC 1) of the hotspot proteins onto the hotspot genotypes of the DO mice (Figure 9). Each hotspot had a distinct pattern of founder allele effects. For the hotspot on chromosome 3, the effects of the B6 and PWK alleles exhibit opposing trends with age (Figure 9A). For chromosome 5, the effect of the AJ allele on protein abundance shows a pronounced trend with age that is not seen for the other founder alleles (Figure 9B). At the chromosome 12 hotspot, the 129 and NOD alleles have larger effects with increasing age but in opposite directions (Figure 9C).

**Figure 9.**
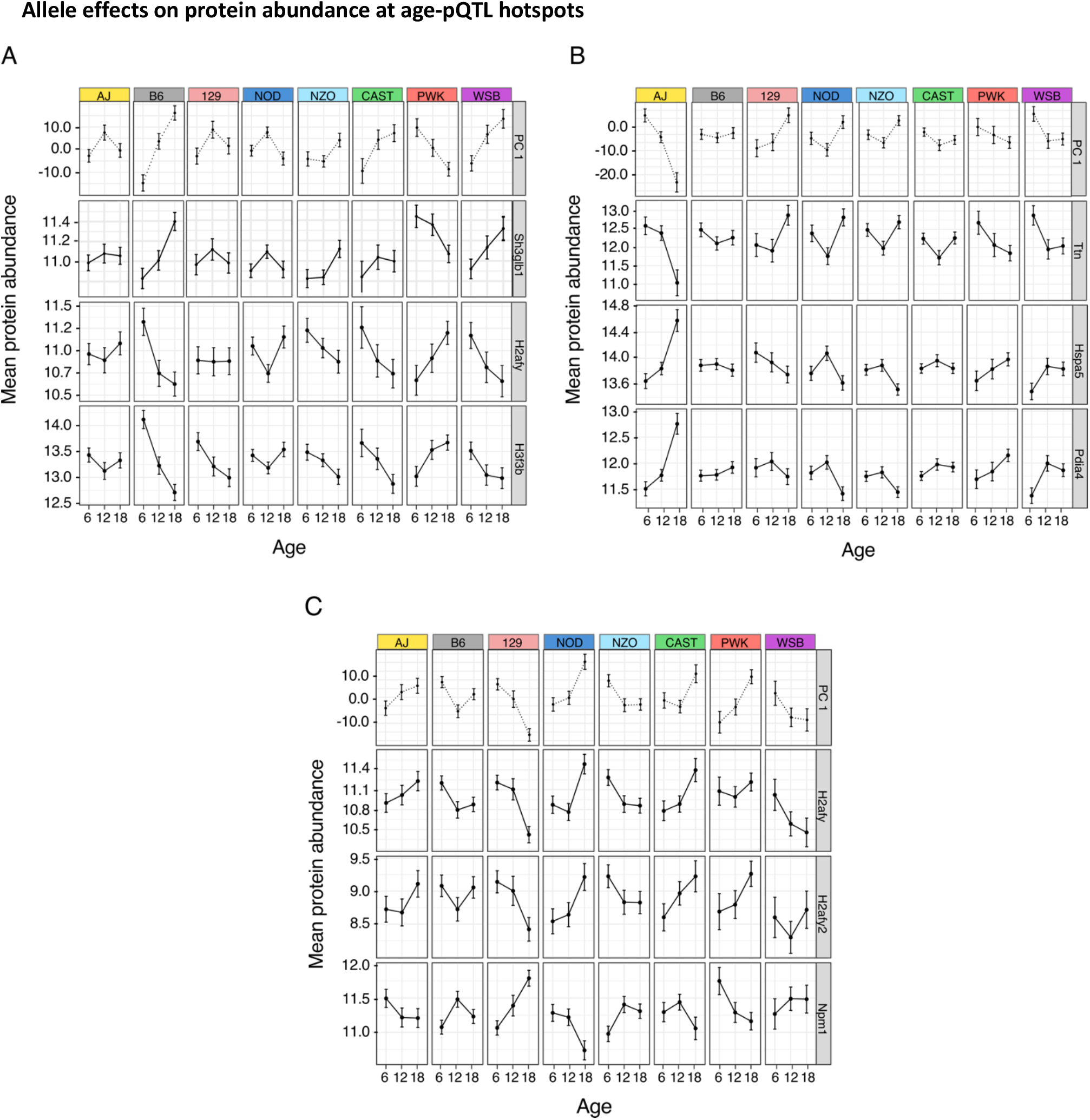
Allele effects on protein abundance at age-pQTL hotspots. Protein abundances associated with DO founder alleles at age-QTL hotspots were estimated for each age group. Estimates are shown for the first principal component (PC 1) of all proteins in the hotspot and for selected individual proteins. Error bars denote +/- 1 SE. Protein SH3GLB1, also known as BIF-1, is located near the hotspot on chromosome 3, and it is positively correlated to PC 1 (A). Proteins H2AFY and H3F3B, in the same hotspot, participate in the nucleosome complex, and are negatively correlated with PC 1, showing inverted allele-specific patterns of change with age (A). HSPA5 and PDIA4, in the chromosome 5 hotspot, participate in the endoplasmic reticulum stress response. These proteins are negatively correlated to PC 1 and show similar allele effect patterns (B). Another chromosome 5 hotspot protein, TTN, which is involved in sarcomere structure, shows opposite allele effects with age (B). H2AFY and H2AFY2, associated with chromatin structuring, mapped to the chromosome 12 hotspot, are positively correlated to PC 1, and share similar allele effect patterns (C). The protein NPM1, also involved in chromatin structure as a histone chaperone, shows inverted allele effects with age (C).

We confirmed that the allele effects for individual hotspot proteins correspond to the estimates based on PC1 but note that some proteins are negatively correlated with PC1 and have opposite direction of change with age (Figure 9). The chromosome 3 hotspot includes members of the histone families (H2AFY and H3F3B), that participate in chromatin structuring and are positively correlated (Supplementary File 4). For these proteins, B6, CAST or WSB alleles drive decreasing abundance with age, while the PWK allele is associated with increasing abundance with age (Figure 9A).

Chromosome 5 hotspot proteins HSPA5 and PDIA4 participate in the physiological endoplasmic reticulum (ER) stress response and are positively correlated and, for mice with the AJ allele, increases with age (Supplementary File 4). Whereas TTN, a protein responsible for sarcomere structuring and muscle contraction, is negatively correlated and, for mice with the AJ allele, TTN abundance decreases with age (Figure 9B).

Proteins related to chromatin structure, including H2AFY, H2AFY2 and NPM1 are found in the chromosome 12 hotspot. Interestingly, H2AFY also mapped to the chromosome 3 hotpsot. The histone proteins, H2AFY and H2AFY2, are positively correlated and, for mice carrying the 129, decrease in abundance with age. The protein NPM1, which encodes a histone chaperone, is negatively correlated to both H2AFY and H2AFY2, and shows opposite allele effects (Figure 9C).

The QTL hotspots identify loci where genetic variation that presumably acts on gene(s) local to the hotpot, influences multiple proteins with distant coding genes. Therefore, we looked for genes within the QTL support intervals of the hotspots that could be the genetic drivers. Contiguous to the hotspot on chromosome 3 (∼145 – 150Mb), we identified a suggestive local pQTL (LOD_int_ = 7.01) for the protein SH3GLB1, also known as BIF-1. The gene is located between 144.68 – 144.72Mb and has an age-by-genotype allele effects pattern similar to the PC 1 allele effects (Figure 9A), consistent with a shared genetic driver and, given the genomic location of SH3GLB1, consistent with the hypothesis that SH3GLB1 modulates the proteins that mapped to the chromosome 3 hotspot. We note that BIF-1 plays a role in autophagy regulation (77,78), an important component of protein quality control system that declines with age (79). We propose BIF-1 as positional candidate driver of the age-related changes in proteins that mapped to the chromosome 3 hotspot. Examination of the chromosome 5 and chromosome 12 hotspot regions did not identify any strong candidate genes.

## DISCUSSION

By analyzing both transcripts and proteins in the mouse heart we were able to detect distinct biological processes that are altered through the course of natural aging. We and others have found that age-related transcriptional changes are not necessarily followed by a change in their proteins and likewise, age-related changes in proteins are not preceded by transcriptional changes (12,13,62,80). We found that transcripts decreasing in expression with age are associated with cardiomyocyte contraction and survival, and fatty-acid metabolism. These findings are consistent with previous reports describing disrupted Ca2+ handling, progressive loss of myocytes, and metabolism switch in the aging heart, which may reflect a compensatory mechanism to improve contractility (3,4,81). Transcripts increasing in expression with age are involved in the acute-phase response and cardiac fibrosis, which suggest chronic inflammation and loss of cardiomyocytes that stimulate extracellular matrix remodeling during the aging process (82–84).

Proteomics reveals major features of the aging heart that are not detected at the transcript level. We saw changes in mitochondrial respiratory chain proteins, as well as intracellular protein and vesicular transport pathways, in agreement with previous proteome analysis of the aging mouse heart (85,86). We observed that most of the proteins from the RAB and AKT families increase their abundance with age, suggesting growth of cardiomyocytes and increased cellular synthesis, which is commonly observed in hypertrophic hearts (1). Proteins of the mitochondrial respiratory complexes II, III and IV increase with age while proteins from Complex I change in both directions. This increase in proteins across most of the mitochondrial complexes could act as a compensatory mechanism in response to an input deficiency from early steps in oxidative phosphorylation.

We found that ACTN3, which is involved in the structuring of sarcomeric Z line and the regulation of the contraction apparatus, decreases with age. *Actn3* expression has been observed in the skeletal muscle, but has not been reported previously in the heart of either mice or humans (87). In knockout mice, the deficiency of ACTN3 in the skeletal muscle promotes a switch from fast-anaerobic metabolism towards a more efficient aerobic metabolism, leading to better exercise performance (58,59). The same study also speculated that the *Actn3* null allele has been under positive selection in modern humans, suggesting that aerobic metabolism boosting may confer a fitness advantage (59). In humans, ACTN2 and ACTN3 appear to have functional redundancy, and *Actn2* was recently linked to heart failure in a multi-omic analysis (87,88). Our data suggests that ACTN3 may participate in the regulation of metabolic efficiency not only in the skeletal muscle, but also in the mouse heart, and might contribute to the physiological changes in muscle efficiency during the aging process.

We observed that proteins associated with fetal metabolism increase with age. A return to embryo/fetal gene program occurs in some pathological heart conditions and it is thought to play a protective role (89,90). The fetal heart is under constant stress conditions, including hemo-dynamic load and hypoxia, and uses carbohydrates as a primary energy source to maintain cardiac efficiency (89,91). The aging heart undergoes different types of stress including the accumulation of reactive oxygen species, cardiomyocyte loss, mitochondrial dysfunction and hypoxia (81,92). These stressors are even more pronounced when they co-occur with other age-related comorbidities, such as diabetes mellitus and hypertension. Studies have demonstrated a return to the fetal gene program in diabetic rats with cardiac diseases (93,94). We propose that a return to the fetal program happens in mice during the normal aging process and may play an adaptive role to compensate for age-related decline in other biological functions of the heart.

The protein data suggest age-related changes in the stoichiometry between members of protein complexes. In general, the correlations between proteins within complexes decrease with age. In contrast, correlations at the transcript level tended to increase with age, although these effects were less pronounced. This dynamic suggests that stochiometric regulation in protein complexes is largely post-transcriptional. These observations are consistent with previous studies that have shown that the production of protein complex subunits is not always perfectly stoichiometric and post-translational mechanisms play an essential role in removing extra subunits and promoting the balance of these complexes (61). Here, in concordance with previous findings using different organisms and tissues (62,63), we propose an age-related loss of stoichiometry across multiple protein complexes, including the mitochondrial respiratory chain complexes, and complexes related to the protein quality control system, notably including the 26S proteasome. Taken together, these findings suggest that age-related protein complexes accumulate stoichiometric imbalance during the aging process.

The proteasome plays a crucial role in regulating the abundance and stoichiometry of proteins in the cell (95), and several studies have attempted to understand why proteasome activity declines with age (96–98). One hypothesis is that reduced expression of proteasome subunits can lead to the decline of proteasome activity in the heart, however, disruption in protein degradation was already observed without reduction in proteasome abundance (98). In this work, we did not observe a reduction in the expression of proteasome subunits, in fact, most of them increase with age (Supplementary File 1; Supplementary File 2). It also has been proposed that post-translational modifications and the excess of oxidized proteins contribute to the decline in proteasome activity in the heart (98,99). Here, supported by previous findings (62,63), we propose that age disrupts the balance of proteasome subunits in the heart, which potentially contributes to a vicious cycle of progressive protein quality control breakdown during the aging process.

For the first time, we found that genetic variation can modulate the dynamics of proteins involved in protein quality control. Most age-QTLs are distant, indicating that genetic variation is not primarily acting on proximal genes, which supports the importance of post-transcriptional mechanisms in regulating protein homeostasis during aging (11,13). Proteins that map to the hotspot loci are functionally related and many have established roles in aging.

Proteins that map to hotspots on chromosomes 3 and 5 are enriched for functions in the proteasome complex and endoplasmic reticulum (ER) lumen, respectively (Figure 8A/B). Proteosome complex proteins PSMD13 and PSMD14 that were also implicated in the loss of stoichiometry (Figure 6D2). These findings suggest that genetic variation can potentially affect the stoichiometry of the proteasome complex. Both the ER and the proteasome complex are part of the protein quality control system that coordinates proteostasis and cell survival (100). The ER contains transmembrane proteins that sense the presence of misfolded proteins, which then activates transcription factors that up-regulate the expression of ER stress response genes (100). Some misfolded proteins are marked by ubiquitination and transferred to the cytosol, where they are degraded by the proteasome complex (100). The proteins HSPA5, HSP90B1 and PDIA4, present in the chromosome 5 hotspot (Figure 8B), are known to be downstream targets of the ATF6 branch of the ER unfolded protein response and this pathway was found to be adaptive and cardioprotective in several studies (101–104).

Enrichment analysis of the hotspots on chromosomes 3 and 12 identified proteins associated with chromatin structure (Figure 8A/C), including histones and histones chaperones, such as NCL and NPM1 (Figure 8C). Loss of chromatin structure was previously reported in aging models for yeast and human fibroblasts (105,106), and leads to dysregulation of global gene expression, increased genomic instability, and promiscuous access of DNA damaging agents to the chromatin (107,108). It is interesting to note that all histone proteins in our data decrease with age (Supplementary File 2), suggesting that age-related loss of chromatin structure also occurs in the heart.

We identified the protein SH3GLB1 (BIF-1) as a potential driver of the age-related dynamics of proteins that mapped at the chromosome 3 hotspot. SH3GLB1 is a member of endophilin protein family that plays a role in mitochondrial fission, vesicle formation and autophagy (77,109,110). As mentioned earlier, the protein quality control system contributes to the aging process and studies suggest that increased autophagy may be a compensatory effect when the proteasome complex is disrupted (111,112). In addition, autophagy plays an important role in maintaining chromosomal stability, and loss of *Sh3glb1* was shown to induce DNA damage due to metabolic stress (113). Autophagy is crucial for maintaining cardiac function - mice with disrupted autophagy show anomalies in sarcomere and mitochondria structure (114). Our findings suggest that genetic variation in the DO mice, near the *Sh3glb1* locus, influences individual rates of change with age of proteins involved in the proteasome complex, chromatin structuring and muscle apparatus organization.

In summary, we have described changes that occur with normal aging in heart tissue from DO mice that suggest a scenario of mitochondrial dysfunction, physiological hypertrophy, and a return to the fetal gene expression program. The proteome data revealed aspects of aging in the heart that are not seen at the transcript level, including the loss of stoichiometry within protein complexes involved in protein trafficking and sorting. We found that genetic variation can influence the age-related dynamics of members of the protein quality control system and other biological processes that play a role in aging. These genetic effects are distal and complex, likely acting through a gene or set of genes near the hotspot locus that indirectly modulates the expression of functionally related groups of proteins. Our findings illustrate how transcriptome and proteome profiling data collected in a genetically diverse model system can reveal broad patterns of change in the molecular dynamics of the aging heart.

## MATERIAL AND METHODS

### Study cohort and tissue collection

The cross-sectional aging study was initiated with 600 DO mice (300 of each sex) bred at The Jackson Laboratory (stock no. 009376) across five waves representing breeding generations 8 through 12 of the DO stock. Mice were maintained on a standard rodent chow (LabDiet 5K52, St. Louis, MO) in an animal room that was free of pathogens, had a set temperature ranging from 20-22° C, and a 12-hour light/dark cycle. The full study was populated across 6 generations of DO breeding over a period of two years. Animals were housed at 4 mice per pen and pens were randomly assigned to 6, 12, and 18 month-aged groups for tissue collection. Whole hearts were dissected, flash-frozen, pulverized and aliquoted. A subset of 192 samples, balanced across age groups and sexes were selected for RNA-seq and shotgun mass spectrometry. This set included 34 females and 30 males at age 6 months, 31 females and males at age 12 months, and 31 females and 35 males at age 18 months. The representation of generational waves included 43 mice from generation 8, 35 mice from generation 9, 38 mice from generation 10, 39 mice from generation 11, and 37 mice from generation 12. The mass spectrometry was performed for 190 of the 192 mice (1 female and 1 male in the 12 months were not included). The Jackson Laboratory Institutional Animal Care and Use Committee approved all procedures used in this study.

### Sample size determination

The sample size required to detect a significant age effect is determined by the expected size of the effect (difference in means between the 6- and 18-month age groups) relative to the within age-group variance. Therefore, we define the strength of an effect in units of standard deviation (SD) of the within group variance. Based on standard power calculations (115), with a sample size of ∼64 animals per age group we can expect to achieve power = 0.80 to detect an age effect of 0.5SD at an unadjusted type I error of 0.05. In practice, because of the high precision of the RNA and protein quantification, this enabled us to detect age effects in the majority of genes tested even after applying a false discovery error rate correction for multiple testing (116). To evaluate the power for genetic mapping, we referred to simulations conducted by Gatti et al. (2014) (117). After applying family-wise error rate for multiple testing of the genome-scan, our sample of 188 animals has expected power = 0.80 to detect a QTL that explain 20% of the total variation in RNA or protein expression.

### Bulk RNA extraction

The frozen and pulverized heart tissue was lysed in Ambion TRIzol reagent (Thermo Fisher Scientific #15596026, Waltham, Massachusetts). Bulk RNA was isolated using the miRNeasy Mini kit (Qiagen Inc. #217004, Germantown, MD), according to the manufacture’s protocols with the DNase digest step. RNA concentration and quality ratios were assessed using the Nanodrop 2000 spectrophotometer (Thermo Fisher Scientific) and RNA 600 Nano LabChip assay (Aligent Technologies, Santa Clara, CA).

### RNA sequencing and quantification

Poly(A) RNA-seq libraries were generated using the TruSeq Stranded mRNA Library Prep Kit (Illumina, San Diego, CA). Libraries were pooled and sequenced 100 bp single-end on the HiSeq 2500 (Illumina) using TruSeq SBS Kit v4 reagents (Illumina). The RNA-seq experiment was performed in two replicates for each sample distributed across 8 lanes. The replicates of each sample were carried out in different lanes to avoid lane effects.

Expectation-Maximization algorithm for Allele Specific Expression (EMASE) was used to quantify multi-parent allele-specific and total expression from RNA-seq data (118) using the Genotype by RNA-seq (GBRS) software package (https://gbrs.readthedocs.io/en/latest/) (RRID:SCR_020963). Transcripts were removed if they did not have at least one read in at least half of the samples, resulting in a total of 20,932 transcripts for further analysis. RNA-Seq counts were normalized relative to total read counts using the variance stabilizing transform (VST) as implemented in DESeq2 (RRID:SCR_015687) (28). For the genetic mapping analysis only, in order to minimize the impact of outliers, we transformed the vst normalized data to rank normal scores (119).

### Mass spectrometry and protein quantification

Mass spectrometry (MS) experiment and protein quantification were performed as described in Chick et al. (2016) (69). In summary, tissue from the total heart samples were homogenized in 1 ml lysis buffer, which consisted of 1% SDS, 50 mM Tris, pH 8.8 and Roche complete protease inhibitor cocktail (Roche # 11697498001, Clifton, NJ). Peptide measurements were performed using Tandem Mass Tags (TMT) and carried out in batches of 10 samples each. MS spectra assignments were made using the Sequest algorithm (120) with the Ensembl database (mouse: Mus_musculus NCBIM37.61), and protein abundances were estimated from their component peptides. Samples were assigned to batches in randomized order. Sample labels were not masked and there were no technical replications performed.

Prior to protein abundance estimation, we filtered out peptides that contained polymorphisms in DO mice relative to the mouse reference genome in order to minimize false pQTL signals that can occur due to failure to detect polymorphic peptides. To estimate and normalize protein abundance from component peptides, we followed Huttlin et al. (2010) (121) and calculated:

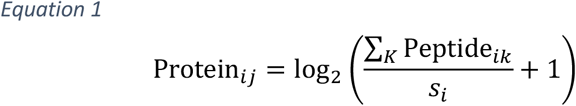

where *K* represents the set of observed peptides that map to protein *j* for mouse *i* and *s*_*i*_ is a scaling factor for standardizing samples within a batch. 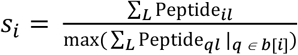, where *L* is the set of all peptides observed for a sample, *b*[*i*] denotes the batch of sample *i*, and max(∑_*L*_ Peptide_*ql*_ : *Q ∈ b*[*i*]) is the maximum sum of peptides for all the samples in batch *b*[*i*]. Protein abundance levels that were missing (NA) were imputed to be zero. Proteins with zeros for more than half the samples were excluded, resulting in a total of 4,062 proteins for further analysis.

The MS experiment was performed in batches of 10 samples processed and quantified simultaneously. Batch effects were removed using a linear mixed effect model (LMM) fit with the lme4 package (RRID:SCR_015654) (122). The batch effect, estimated as a best linear unbiased predictor (BLUP), was subtracted from each protein abundance, while age (as a categorical variable with three levels) and sex were included as fixed effect covariates in the model. For genetic mapping analysis, protein abundances were transformed to rank normal scores to minimize the effect of outliers (119).

### Age effect on transcript expression

We used the DESeq2 package (28) to test for transcripts whose expression changed with age. Briefly, we fit the following GLM using the negative binomial distribution in DESeq2:

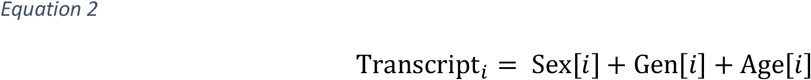

where Transcript_*i*_ is the total count for each transcript from mouse *i*, Sex[*i*] is the effect corresponding to the sex of mouse *i*, Gen[*i*] is the effect corresponding to the generation of mouse *i*, and Age[*i*] is the effect corresponding to the age of mouse *i*, fit as a continuous variable at the year scale (0.5, 1, 1.5 years) obtained by dividing the age groups by 12. The age effect was tested using its Wald statistic and it estimates the log_2_ fold change per year of life. Transcripts with a significant age effect on expression were determined after FDR adjustment to account for multiple testing across all transcripts (FDR < 0.1).

### Age effect on protein abundance

To detect proteins with age effects, we fit a log-normal linear model with predictors similar to Equation 2:

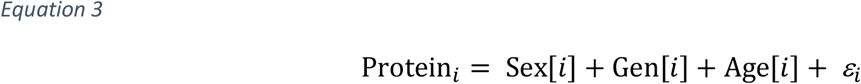

where Protein_*i*_ is the log-scale abundance of each protein from mouse *i*, as defined in Equation 1, *ε*_*i*_ is the residual, and all other terms as previously defined. The age effect corresponds to the slope of the regression model and it estimates the log_2_ fold change per year of life. Proteins with significant age effects were identified after FDR adjustment (FDR < 0.05).

### Age effect in the correlation of genes coding for protein complexes

To investigate the stoichiometry of protein complexes (64) (CORUM database) we adapted a method described in McKenzie et al (2016) (65). We computed the Pearson correlation between the expression of each gene-pair in the protein complexes for both transcript and protein data and for each age group. Then for each protein complex and data type (transcript and protein), we regressed the correlation coefficients of each gene-pair on age and recorded the slope, which represents the age effect, in units of change in correlation per year, as in Equation 4:

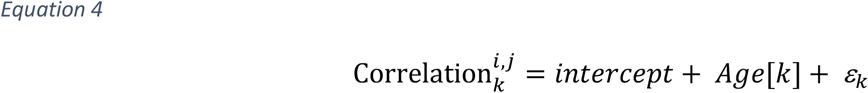

where Correlation 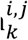 is the correlation between proteins *i* and *j* at age *k, intercept* is the overall intercept, *Age* is the age effect, fit from a continuous encoding of age at the year scale, *Age*[*k*] = *β*_*Age*_ * *k*, and *ε*_*k*_ is the residual at age *k*. In order to determine significance, we shuffled the mouse IDs and repeated the slope estimation 1,000 times to obtain FDR estimates using the DGCA package (RRID:SCR_020964) (65). Significant age effects were declared at FDR < 0.1.

We also fit a model to compute the overall age effect for each protein complex, without fitting separate models per gene-pair. We fit a linear mixed model using the R/lme4 package to jointly model gene-pairs with a random effect, allowing the intercept and age slope for each gene-pair:

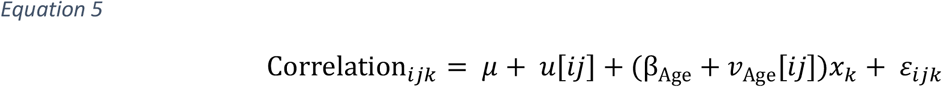

where Correlation_*ijk*_ is the correlation between proteins *i* and *j* at age *k, μ* is the overall intercept, *u*[*ij*] is the random deviation on the intercept specific to the pairing of proteins *i* and *j*, β_Age_ is the overall age effect, *v*_Age_[*ij*] is the random deviation on the age effect specific to the paring of proteins *i* and *j, x*_*k*_ is the age (6, 12, or 18 months), and *ε*_*ijk*_ is random noise on the correlation for proteins *i* and *j* at age *x*_*k*_. The protein pair-specific random terms are modeled as **u ∼** N(**0, I***τ*^2^) and 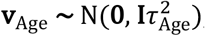, and the error as *ε*_*ijk*_ **∼** N(**0, I***σ*^2^). We used the permutations procedure from McKenzie et al (2016) (65) to determine significance, using a *p*-value cut-off of 0.05, for each protein complex.

### Additive QTL mapping

Although the results of additive QTL mapping are not reported here, it is useful to describe the methods used as a prelude to the description of age-interactive QTL mapping analysis. For each transcript or protein, we transformed the data to rank-normal scores (119) and fit the following model at ∼64,000 equally spaced loci across the genome:

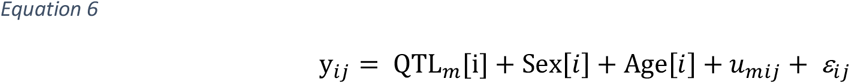

where y_*ij*_ is the transcript or protein *j* expression/abundance for mouse *i*, QTL_*m*=_[*i*] is the expected dosage of founder haplotype alleles for mouse *i* at locus *m, u*_*mij*_ is a random kinship effect that accounts for the correlation between individual DO mice due to shared genetic effects excluding the chromosome of locus *m*. The kinship effect is modeled as 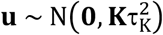, where **K** is a realized genomic relationship matrix and 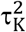 is the variance component underlying the kinship effect (123). The log_10_ likelihood ratio (LOD score) was determined by comparing the QTL model (Equation 4) to the null model without the QTL term.

### Age-interactive QTL mapping

We performed a second set of genome scans to identify age-interactive QTL loci where the rate of change of a transcript or protein is dependent on genotype. Genome scans for age-QTL are based on the following model:

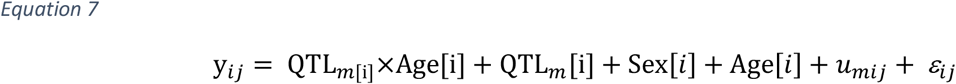

where QTL_*m*[*i*]_×Age[*i*] is the interaction effect between the QTL genotype and age of mouse *i*. All other terms are as previously defined. The null model for the age-interactive genome scans is the model from Equation 6, thus only the interaction term is being tested. To determine significance thresholds for age-QTL we required a more elaborate permutation procedure than the standard used for additive QTL (124). For each transcript or protein, we fit the following model:

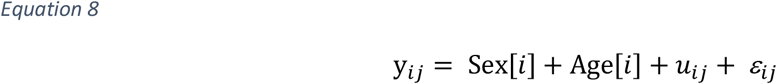

Where the kinship term u_ij_ includes effects of all loci, including the additive effect of the locus under evaluation. We then computed the residuals by subtracting the fitted values of model predictors:

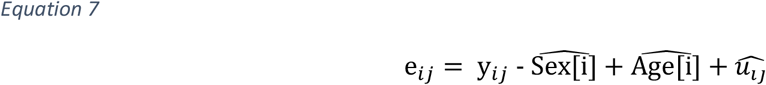

To construct a permutation test, we generate null data by summing the fitted effect values with a randomly permutated of the residuals from Equation 7. We repeated the age-interactive scans on the residual-permuted phenotypes 1000 times to obtain a null distribution sample of the LOD_int_ statistic. Signficance thresholds for the maximum LOD_int_ scores were based on the 95th percentile of this distribution and a suggestive threshold was determined using the 37^th^ percentile. Transcript and protein age-QTLs were considered significant when LOD_int_ > 7.75 and suggestive when LOD_int_ > 6.0. All QTL analyses were performed with the R/qtl2 package (RRID:SCR_020965) (125).

### Distal QTL hotspot analysis

Using a sliding window of 4 Mb, we counted the density of suggestive age-QTL (LOD_int_ > 6) for transcripts and proteins (Figure 7). We defined a genomic region as a hotspot based on having more than 40 age-QTLs. We used the hotspots to defines sets of transcripts and proteins that mapped to these regions. We further refined the hotspot sets by filtering out transcripts or proteins with a mean Pearson correlation coefficient < 0.3 with the other hotspot members, removing genes that are not highly correlated with other genes that map to the hotspot.

### Age-specific allele effects

Proteins with shared genetic drivers, which could cause distal age-QTL hotspots, should have similar allele effects. To investigate this, we computed the principal components (PCs) from the refined hotspot proteins. We then estimated age-specific allele effects for the first principle component (PC1), and for each protein that mapped to the hotspot, using the function fit1() from R/qtl2 package (125). We fit the model separately to each age group, including sex as an additive covariate, to estimate the age-specific effects associated with each of the eight founder alleles that are present in DO mice.

### Functional enrichment analysis

We performed functional enrichment analysis for transcript and protein gene sets with significant age effects and also to sets of genes in the refined age-QTL hotspots. We used the ClusterProfiler package (RRID:SCR_016884) (26) to identify enriched gene ontology terms (biological processes, cellular compartments and molecular functions) for each set. We used stringent (FDR < 0.05) and lenient (FDR < 0.1) cut-offs to defined enriched categories for reporting.

### Software

All data analysis and figures were generated using R v3.6.0 (RRID:SCR_001905) based on the packages tidyverse v1.3.0 (RRID:SCR_019186) and ggplot2 v3.3.0 (RRID:SCR_014601). The R Scripts used for all the analysis performed on this work can be found on Github (https://github.com/isabelagyuricza/Aging_Heart_DO_analysis) and on Figshare (DOI: 10.6084/m9.figshare.12430094.v1).

### Data access

The RNA fastq files can be found on on NCBI SRA repository under bioproject PRJNA510989. The mass spectrometry proteomics data have been deposited to the ProteomeXchange Consortium via the PRIDE partner repository (http://www.proteomexchange.org/-PXD023724). Both raw and normalized transcript expression matrices, as well as the protein abundance data have been deposited to Figshare (10.6084/m9.figshare.12378077). Genotype data is deposited the DODb database (https://dodb.jax.org/). All data and QTL results are available for download and interactive analysis using the QTLViewer for user driven queries (https://qtlviewer.jax.org). Raw and normalized quantification of transcripts and proteins and the genotype data are included in the download in a Rdata format suitable for further analysis.

## Acknowledgements

The authors gratefully acknowledge the contribution of Matthew Vincent for creating the QTLViewer web tool. We also thank Heidi Munger and all the staff from the Genome Technology Service at The Jackson Laboratory for expert assistance with the RNA-seq experiments described in this publication. This work was supported by the National Institutes of Health and The Jackson Laboratory Nathan Shock Center of Excellence in the Basic Biology of Aging (P30 AG038070).

## Supplementary material legends

**Figure 5 – figure supplement 1**. Standardized age effects for the expression/abundance of all the subunits from the mitochondrial complexes (I-V) at the transcript (x-axis) and protein (y-axis) levels. Color indicates the significance of the age effect at the transcript level (FDR < 0.1), protein level (FDR < 0.05), both or neither. For all the complexes, significant age-related changes are predominant at the protein level, and, with exception of mitochondrial complex V, they tend to change in both directions. Age effect corresponds to the linear trend with age in units of LFC per year.

**Figure 7 – figure supplement 1**. Heatmaps for the Pearson correlation coefficients between transcripts or proteins that mapped to the age-QTL hotspots. A) Correlations between transcripts that mapped to the chromosome 4 age-eQTL hotspot show that there is no evidence for correlation in this group of transcripts. B) Correlations between proteins that mapped to the chromosomes 3 (B1), 5 (B2) and 12 (B3) age-pQTLs hotspots identify tightly correlated groups of proteins (positive or negative) that are potentially regulated by shared genetic variation.

**Supplementary File 1**. List of transcripts with significant age-related changes in expression.

**Supplementary File 2**. List of proteins with significant age-related changes in abundance.

**Supplementary File 3**. List of gene-pairs members of protein complexes that have a significant change in their pairwise correlations with age at the transcript or at the protein level.

**Supplementary File 4**. Annotation of all the suggestive age-eQTLs and age-pQTLs.

