## Supplementary figures and images for "Genome-wide transcript and protein analysis reveals distinct features of aging in the mouse heart"

### Figure 5 figure supplement 1

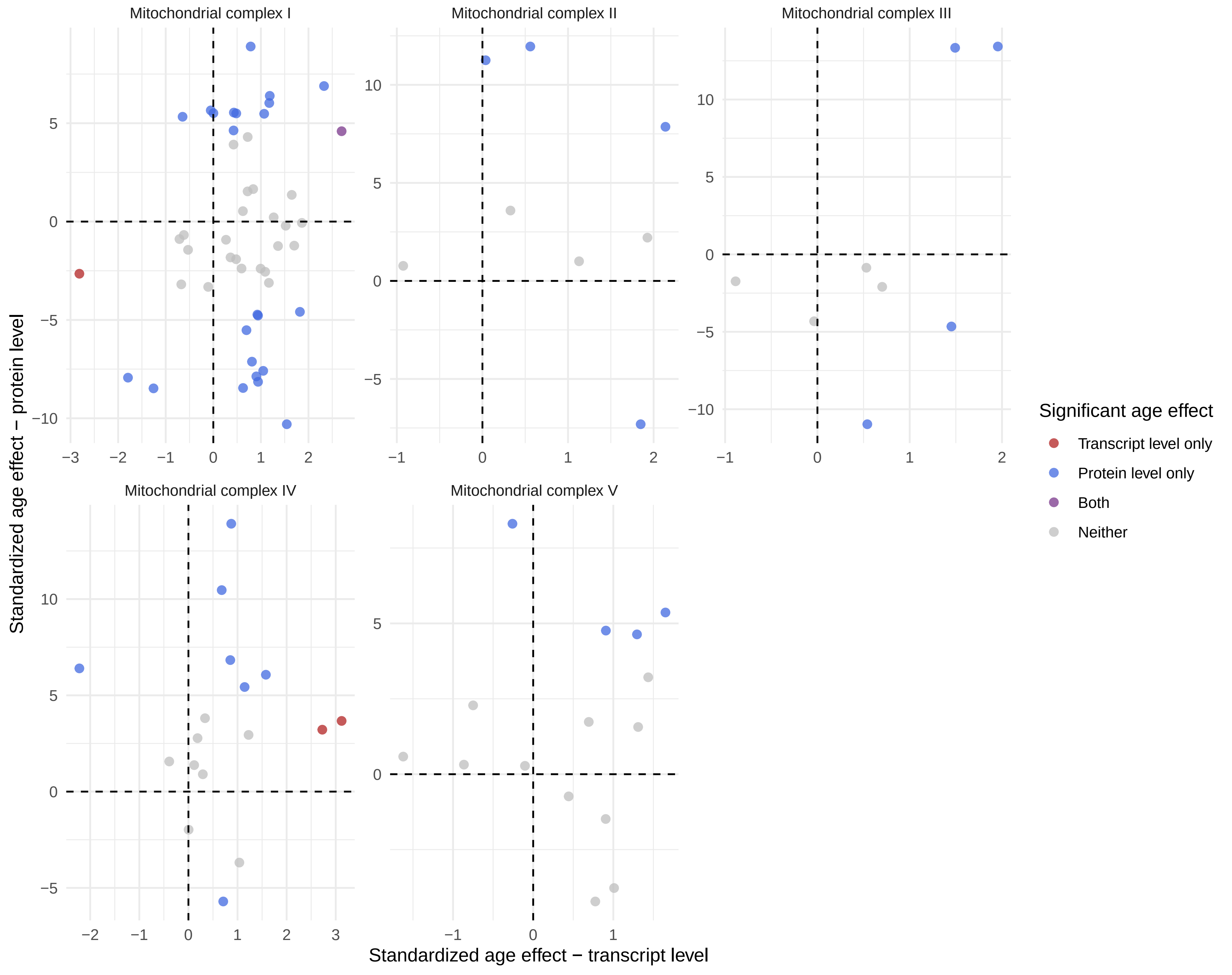

### Figure 7 figure supplement 1

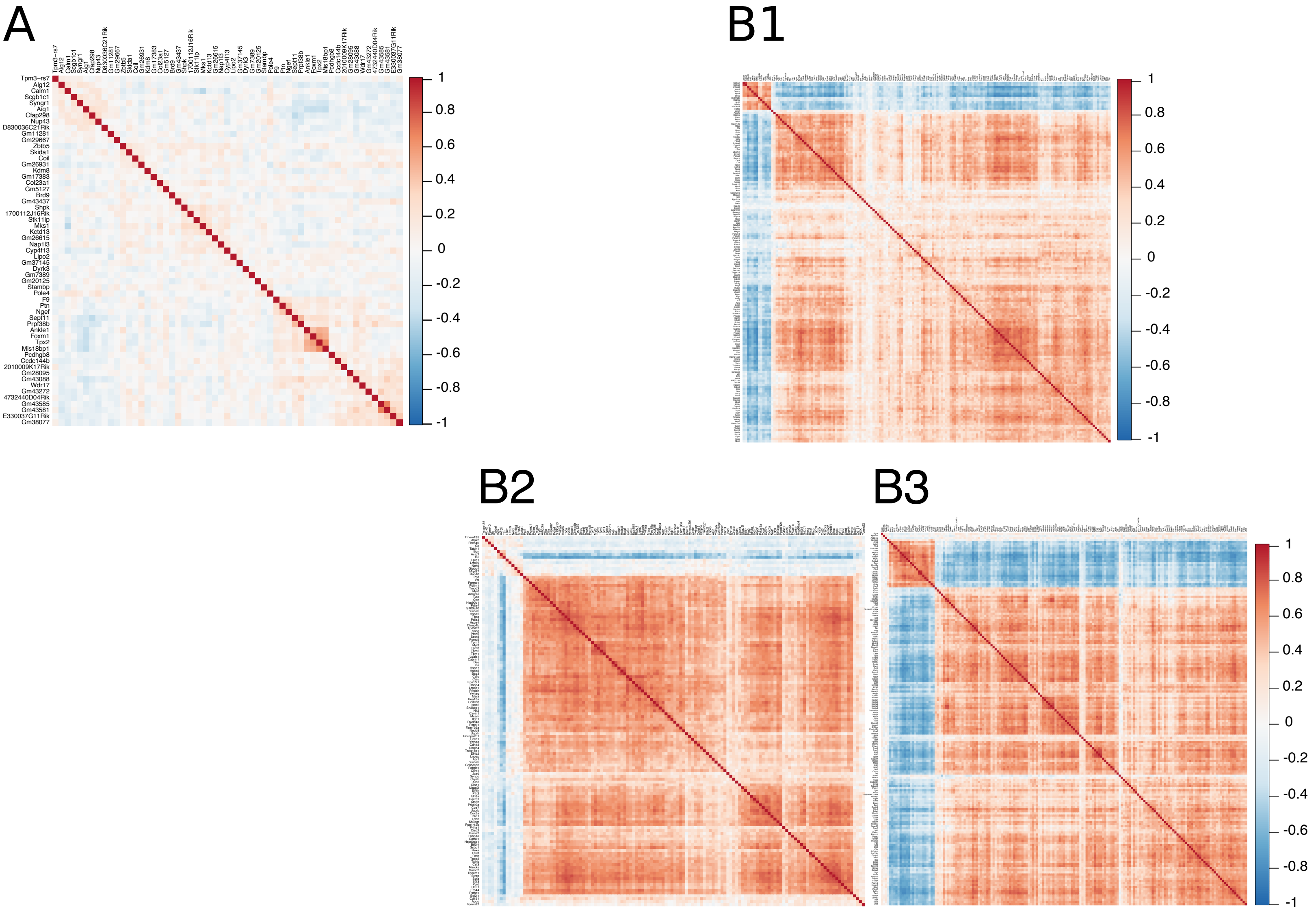
